# Coexpression enhances cross-species integration of scRNA-seq across diverse plant species

**DOI:** 10.1101/2023.11.28.569145

**Authors:** Michael John Passalacqua, Jesse Gillis

## Abstract

Single-cell RNA sequencing is increasingly used to investigate cross-species differences driven by gene expression and cell-type composition in plants. However, the frequent expansion of plant gene families due to whole genome duplications makes identification of one-to-one orthologs difficult, complicating integration. Here, we demonstrate that coexpression can be used to identify non-orthologous gene pairs with proxy expression profiles, improving the performance of traditional integration methods and reducing barriers to integration across a diverse array of plant species.

## Main

Plants have a remarkably flexible cellular physiology, driving their adaptation into nearly every environment. Recently, the advent of single-cell RNA sequencing (scRNA-seq) has provided novel insights into the diversity of cell types underlying these adaptations^1,2^. The unique diversity in plants makes comparative assessments between species important, but it is also complicated by uncertain homology relationships. Unlike in mammals, where homologous genes and structures can be easily identified, plant gene families frequently expand by whole genome duplication (WGD), polyploidization and tandem gene duplication^3–5^. This scarcity of one-to-one gene pairs is a major barrier to defining a common gene space for the integration of single-cell data, a key step for successful cross-species comparative analysis or integration^6,7^. With vast amounts of single-cell plant RNA data becoming available^8^,this study aims to address a critical gap in its analysis by using coexpression to identify pairs of genes that, while not exclusive orthologs, are functionally related enough to enable the integration of this high-dimensional data. By reducing barriers to integration, we prime the field for the discovery of novel, cell-type specific innovations that have been critical to plant adaptation and domestication.

While a given plant sample may have thousands of expressed genes, the expression patterns of these genes are not independent, and are instead organized into the regulatory programs which underlie cell types. This coexpression generates the low dimensional expression space that is foundational to the success of modern single-cell analysis^9^. We hypothesize that genes with highly similar expression profiles between two species can be used as reasonable proxies for integrating cell-type specific data, that we can identify such profiles using coexpression, and that this will expand the shared gene space, improving our ability to compare cross-species data. The essence of the approach is to use meta-analysis from prior bulk mRNAseq data to define cross-species gene pairs (coexpression proxies) that can be applied in more specific, but sparser, single cell data. By utilizing robust coexpression networks built from over 16,000 publicly available RNA sequencing datasets, as well as gene phylogenies from OrthoDB, we ensure that the coexpression proxies accurately reflect the underlying biology of each species pair they are drawn from^10,11^. This use of gene phylogenies builds upon previous work to improve cross-species integration by relaxing homology requirements^12^. We validate the coexpression proxies with two test examples, highlighting their utility. In the first test, we show that coexpression proxies can accurately reintegrate a split dataset with no shared gene space. Second, we show that the coexpression proxies improve the integration of real single-cell data between two species with complex genomes, maize and rice.

Our first test is an extreme one in which we attempt to integrate two datasets with neither shared genes nor direct orthologs. This would be impossible with a traditional integration approach. To construct this as a case with a known integration, we split an Arabidopsis single cell dataset into two halves, each assigned a mutually exclusive half of the genome. This provides a set of cells with known underlying cell types, shared across two datasets, but with no shared genes. To integrate in this otherwise uncorrectable scenario, we used coexpression proxies from bulk data to identify within genome pairs. As an example, the selected coexpression proxy gene, AT1G16150, closely matches the expression profile of the target gene, AT1G16160. In contrast, AT4G31100, a rejected gene from the same ortholog family, has a distinct expression profile (Figure 1B). We quantify this on a per-gene basis by assessing the degree to which rejected and selected gene pairs show the same expression across cell-types (measured as by Euclidean distance). We find that accepted coexpression proxies are much closer to the target’s expression profile across cell types, and that the rejected proxies are on average 83% further from the target’s expression (Figure 1F). This shows that the coexpression proxies are more similar in expression profile to their target genes than even other genes from the same orthogroup.

**Figure 1.**
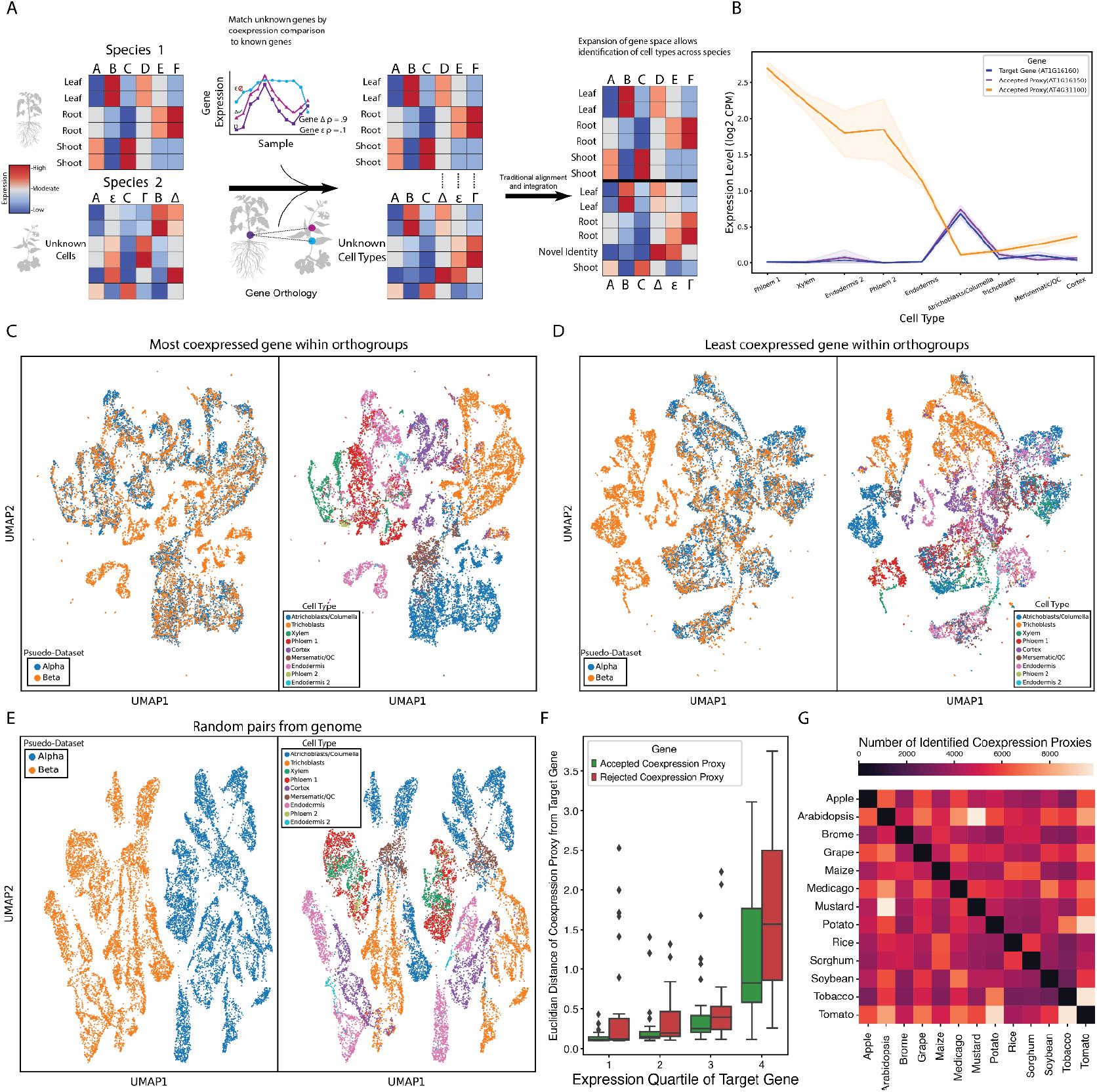
Coexpression proxies integrate a split dataset without shared genes. **A)** Schematic: identification of coexpression proxies improves the integration of single cell data. **B)** Gene expression profile for target gene (AT1G16150) and two potential coexpression proxies (AT1G16160, AT4G31100). The gene with the more similar profile, AT1G16160, was identified as a coexpression proxy, while AT4G31100 was rejected. **C)** UMAP showing integration of a split and disassociated *Arabidopsis thaliana* dataset using coexpression proxies. **D)** UMAP showing integration of the same dataset using the worst potential coexpression proxy from each gene family. **E)** UMAP showing the failed integration of the split and dissociated dataset using 1900 random gene pairs. **F)** Euclidian distance from the expression profile of the target gene for select accepted coexpression proxies and rejected coexpression proxies, split by expression quartile of the target gene. **G)** Heatmap showing the number of identified coexpression proxies between each species pair in the database.

Next, we used these coexpression proxies to reintegrate the split Arabidopsis dataset. Using Scanorama^13^, we reintegrated and re-clustered the dataset, placing 82% of cells into a cluster with cells from both datasets (Figure 1C). Additionally, the reintegration was accurate, successfully matching cells of the same cell type across datasets 75% of the time. To evaluate how much of the gene proxies’ success is dependent on information from the gene phylogenies, and how much information is derived from the conservation of coexpression profile, we attempted to integrate the datasets using the worst rejected proxy from within each ortholog group (i.e., lowest coexpression). We find that integration is still possible using these gene pairs, although performance is lower, reducing the successful matching of cells to 65% (Figure 1D). This moderate performance suggests that simple relaxation of orthology constraints is a substantial contributor to performance. However, coexpression provides a significant overall signal boost. This was particularly clear for phloem, which was otherwise unintegrated or mixed with atrichoblasts and xylem. To determine whether sequence similarity alone would prove sufficient, we calculated the pairwise protein sequence similarity of every Arabidopsis gene and attempted to use this to identify gene proxies. While able to perform better than random, this metric was worse than coexpression at reintegrating the split dataset, and completely failed to reintegrate certain clusters. Finally, we attempted integration using 1900 random gene pairs and find that we are unable to achieve any integration (Figure 1E). Given the success of our approach, we generated coexpression proxies between 13 plant species and identified an average of 5,056 gene pairs between species (Figure 1G). The coexpression proxies are numerous enough to provide additional information across even highly diverged species and are well represented (5283 pairs) even between *Zea mays* and *Arabidopsis thaliana*, which diverged 160 MYA. Importantly, while we used Scanorama, these co-expression proxies can be easily incorporated into any potential integration pipeline as they simply expand the shared feature space.

Having shown that coexpression proxies could integrate an otherwise uncorrectable dataset, we tested their ability to improve the integration of single cell data across two different species. Using a supervised integration, we attempted the integration of two root datasets, one from maize and one from rice. We focused on broad cell types for which author annotations directly aligned. Using coexpression proxies, we successfully integrated the maize and rice dataset, accurately integrating 37% of cells into clusters with cells from both datasets (Figure 2A). The remaining cells were different enough to still appear as distinct sub-clusters across species. While this is far from 100%, real cross-species differences do exist, so it is not clear what the maximum plausible percentage is. Importantly, our integration is better than using only the 1-1 gene pairs, which integrated only 14% of the cells (Figure 2B). Key cell-types, such as epidermis and stele, are well integrated using coexpression and missed by 1-1 gene pairs. Similarly, coexpression did not overfit real differences away, capturing the likely real difference between cortex cells, where constitutive aerenchyma formation is critical to oxygen diffusion in partially submerged rice^14^. To evaluate the integration on a cell-type by cell-type basis, we employed MetaNeighbor, which enables us to quantify the degree to which cell types replicate across datasets in a statistical framework ^15,16^. We compare 4 integrations using scGen — utilizing coexpression proxies and 1-1 genes, only coexpression proxies, only 1-1 genes, and using random genes (Figure 2C). As can be seen, coexpression proxies alone, 1-1 pairs alone, and the combination all accurately and similarly group cell types across species. While subtle for this broad classification, the full coexpression proxy set integrates better than either of its parts in all cell types when evaluated by MetaNeighbor (except cortex, where all methods are perfect), reflecting the additional information from the coexpression proxies. Because this is a validation focused on well-defined alignment, performances generally go from high to even higher (e.g., stele goes from AUROC 0.93 to 0.973). To evaluate utility of an increased known gene-pair space, as well as the robustness of the model, we swapped in coexpression proxies for random pairs and tracked performance improvement (Figure 2D). Performance increases steadily to near 1 for most cell types, indicating that the typical number of 5000 coexpression proxies is sufficient to integrate cross-species data. Further querying the coexpression proxies, we found they typically represented core conserved functions such as photosynthesis, mitochondrial proteins, and ribosome metabolism (Figure 2E).

**Figure 2.**
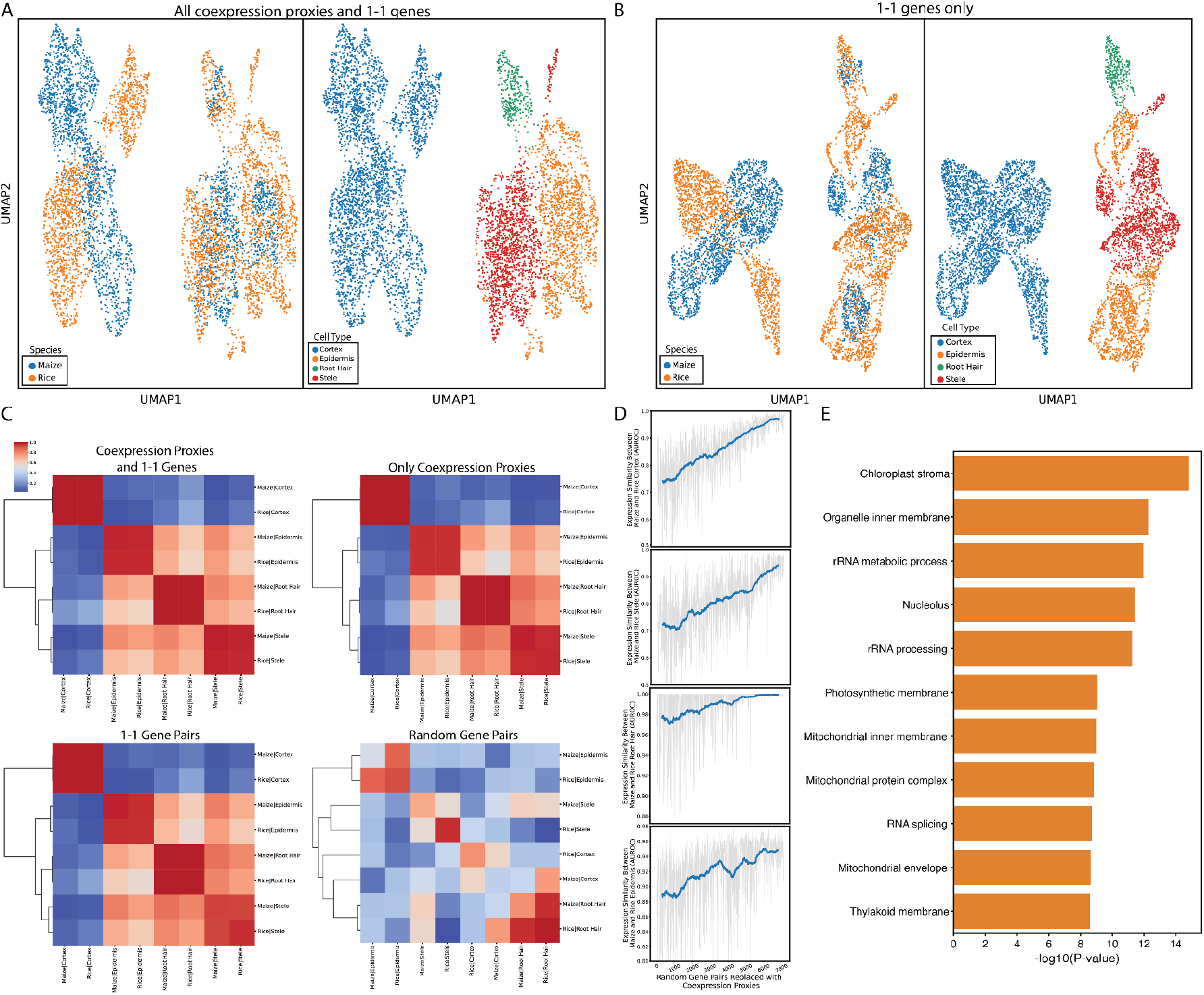
Integration of maize and rice scRNAseq data using coexpression proxies. **A)** UMAP showing integration of *Zea mays* and *Oryza Sativa* using coexpression proxies. **B)** UMAP showing integration of *Zea mays* and *Oryza Sativa* using only 1-1 gene pairs from OrthoDB. **C)** Metaneighbor plots showing post integration similarity between cell types using 4 different gene sets. **D)** Improvement in integration across 800 integration runs as random gene pairs are gradually swapped for coexpression proxies. **E)** Enriched GO terms among rice-maize coexpression proxies

Integrating cross-species single-cell data is an increasingly common goal in the fields of plant development, evolution, and molecular biology. To facilitate this process, we have demonstrated that using coexpression proxies expands the gene space available for integration. To facilitate adoption of this approach by the community, we have generated pairwise coexpression proxies between 13 plant species at 3 thresholds. All coexpression proxy lists are made available at https://gillislab.shinyapps.io/epiphites/. These proxy lists provide an important resource for improving the integration of single-cell data, accelerating the transfer of knowledge from well-studied model organisms to crop systems that are crucial to the global food supply.

## Acknowledgements

We thank Drs. Kenneth Birnbaum and Bruno Guillotin for their discussion on cortex in monocots and their feedback on the manuscript. We thank Drs. David Jackson and Xiaosa Xu for their inspiration of the project and their feedback on the manuscript. We thank John Hover for his support in setting up the website. We acknowledge funding support from NSF IOS-1934388 and NIH R01 MH113005. We acknowledge support from William Randolph Hearst Foundation and the CSHL School for Biological Sciences.

## Methods

### Public gene expression data

All analyses were performed in Python 3.9 and SCANPY^17^. Aggregate coexpression networks were downloaded from CoCoCoNet^10^. Arabidopsis thaliana single-cell RNA-seq expression data from 4 datasets were downloaded from the Gene Expression Omnibus GEO IDs: GSE116614, GSE121619, GSE123818, GSE123013)^1,2,18,19^. Cluster assignments were downloaded from GEO for IDs GSE121619 and GSE123013, or provided by the authors for IDs GSE123981 and GSE116614. *Oryza sativa* single-cell RNA-seq expression data and accompanying cluster assignments were downloaded from GSE146035^20^. *Zea mays* single-cell RNA-seq expression data and accompanying cluster assignments were downloaded from GSE183171^21^, and only nitrate treated cells were used.

### Gene coexpression proxy identification

For each species pair, gene family orthology information was downloaded from OrthoDB V10^11^. Utilizing one to one gene pairs, coexpression conservation was calculated between all genes in each species^22^. Briefly, we compare each gene’s top 10 coexpression partners across species, limiting the potential coexpression partners to genes that are also one-to-one orthologs. We use the ranks from one species to predict the coexpression partners of the second species, and then repeat this in the other direction, averaging the scores to generate the conservation of coexpression score, which is an area under the receiver operator curve (AUROC). This resulted in a Species A genes by Species B genes matrix, filled with the AUROC score for each gene pair. For each gene family, the coexpression conservation matrix was filtered to every possible cross species gene pair. Next, pairs in multigene groups were eliminated by thresholding in two steps. First, any gene pairs with scores below a *quality* threshold were discarded. Second, remaining pairs were required to be reciprocal best hits, and to be higher than other potential options by a *multi-pair* threshold. For genes that were one to one matches, they were only discarded if below a lower *single pair quality* threshold. For the moderate filtering, the *quality, multi-pair* and *single pair junk* thresholds were 0.85, 0.03, and 0.8. For lenient and stringent filtering, the thresholds were 0.8, 0.02, 0.07, and 0.9, 0.035, 0.085, respectively.

### Dataset integration and evaluation

To generate an integration task uncorrectable without a shared gene space, the Arabidopsis dataset was split into two sets of cells. One half of the genome was randomly selected and assigned to the first set of cells, with other data being discarded. The second set of cells were assigned the second half of the genome, and the genes assigned to the first half were discarded. Utilizing the same method as above, coexpression proxies were identified between the two halves of the genome with the moderate threshold. Aligning the two gene spaces using these proxies, we performed integration using the Scanorama Python package^13^. The scanorma.integrate function was used to integrate the two datasets into a shared low dimensional space, and this was plotted using scanpy.tl.umap. For evaluation, we defined a cluster of the same cell type as one containing more that 60% of that cell type, and a mixed cluster as one composed of between 30-70% of either batch.

As the cross-species integration scenario was more challenging, it was integrated utilizing scGEN^23^. Utilizing coexpression proxies between rice and maize at the moderate threshold, the two datasets were aligned. The two datasets were first aligned utilizing coexpression proxies between rice and maize at the moderate threshold. Next, the scGEN model was initialized using scgen.SCGEN, and trained using scgen.model.train, using default parameters. Next, the integration was performed using scgen.model.batch_removal. To evaluate the integration beyond the low dimensional representation, MetaNeighbor was used to compare the post integration similarity of cell types^15^. In order to confirm the model was utilizing coexpression proxies and not relying on training information, the integration was ran 888 times, starting with random gene pairs. Following each run, 8 random pairs were replaced with 8 coexpression proxies, until all were replaced. GO term enrichment was performed using Fisher’s exact test to find terms over-represented in the coexpression proxies, utilizing all genes in the bulk network as the background gene set. Multiple hypothesis correction was performed using Benjamini Hochberg correction.

